# Altered neurocognitive processing of tactile stimuli in patients with Complex Regional Pain Syndrome (CRPS)

**DOI:** 10.1101/146613

**Authors:** Anoop Kuttikat, Valdas Noreika, Srivas Chennu, Nicholas Shenker, Tristan Bekinschtein, Christopher A. Brown

## Abstract

Chronic pain in CRPS has been linked to tactile misperceptions and deficits in somatotopic representation of the affected limb. Here, we identify altered cognitive processing of tactile stimuli in CRPS patients that we propose marks heterogeneity in tactile decision-making mechanisms. In a case-control design, we compared middle and late-latency somatosensory-evoked potentials (SEPs) in response to pseudo-randomised mechanical stimulation of the digits of both hands (including CRPS-affected and non-affected sides) between 13 CRPS patients and 13 matched healthy controls. During a task to discriminate the digit simulated, patients (compared to controls) had significantly lower accuracy and slowed response times but with high between-subject variability. At middle latencies (124-132 ms), tactile processing in patients relative to controls showed decrements in superior parietal lobe and precuneus (that were independent of task demands) but enhanced activity in superior frontal lobe (that were task-dependent). At late latencies, patients showed an augmented P300-like response under task demands that localised to supplementary motor area (SMA). Source activity in SMA correlated with slowed response times, while its scalp representation intriguingly correlated with better functioning of the affected limb, suggesting a compensatory mechanism. Future research should investigate the clinical utility of these putative markers of tactile decision-making mechanisms in CRPS.

**Disclosures:** The research was supported by funding from Cambridge Arthritis Research Endeavour (CARE). The study was sponsored by Cambridge University Hospitals NHS Foundation Trust and supported by its Rheumatology department. The authors report no conflicts of interest.

**Perspective:** We present evidence of altered but highly variable cognitive processing (124 - 268ms latency) in response to mechanical tactile stimuli in patients with CRPS compared to healthy controls. Such mid-to-late latency responses could potentially provide convenient and robust biomarkers of abnormal perceptual decision-making mechanisms in CRPS to aid in clinical detection and treatment.

## Introduction

Complex Regional Pain Syndrome (CRPS) is an unexplained chronic, debilitating pain condition that is characterized by disproportionate pain, swelling, vasomotor, sudomotor, trophic and motor changes. The clinical heterogeneity of the condition points to mechanistic heterogeneity ^24^ and there is a clear and unmet need to better characterise patients in terms of underlying mechanisms to aid early detection and targeted treatment ^20^. Observations of tactile misperceptions in patients with CRPS have prompted investigation of somatosensory neuroplasticity as a putative mechanism in CRPS pain ^28^. However, in one such test requiring patients to identify the digit touched without corresponding visual information, only just less than half of CRPS patients perform poorly on the affected hand ^12^. Despite this variable performance across patients, such metrics (which include lower accuracy in discrimination but also longer response times) improve classification of CRPS patients from patients with limb fracture, even if the patients suffer from lower-limb CRPS and are tested on their hands ^21^. This lack of spatial specificity suggests central mechanisms may influence performance decrements and supports further investigation of such mechanisms.

Decrements in tactile spatial discrimination in CRPS patients ^30^ have been associated with neuroplasticity involving shrinkage of the somatosensory cortical homunculus ^30^. Investigations of this phenomenon have used the high temporal resolution of MEG and EEG, focussing on early latency (<100ms) cortical tactile responses arising from somatotopic cortical representations. However, recent fMRI studies have failed to replicate findings of somatotopic changes in CRPS ^29,43^. Furthermore, there is a lack of research investigating later “cognitive” stages of tactile processing that provide alternative central mechanisms for perceptual disturbance.

We hypothesised that deficits in tactile discrimination in patients with CPRS could be related to aberrant perceptual decision-making mechanisms at late latencies. In particular, we posit a role for mechanisms that are related to context-updating, namely the updating of a cognitive representation (mental model) of the environment. According to computational models of decision-making, notably those positing perceptual decision-making as a form of Bayesian inference, context representations are critical for efficient perceptual decision-making by providing top-down constraints on (noisy or ambiguous) lower-level sensory representations ^1^.

A prominent theory ^31^ links context-updating to the P300, a robust late-latency component occurring ~200–400 ms post-stimulus. EEG research in other clinical contexts has found that P300 marks cognitive dysfunction in chronic headache ^10^, chronic lower back pain ^38^, phantom limb pain ^18^, schizophrenia ^41^, disorders of consciousness ^5^ and dementia ^27^. Sub-components of the P300 with different latencies and scalp distributions have been related to endogenous and exogenous processes. The “P3b” component is thought to reflect context updating processes that are sensitive to task demands (providing a marker of endogenous attentional resource allocation). Thus P300 responses typically correlate with stimulus-response times ^34^, providing a potential marker of perceptual decision-making efficiency. On the other hand, the “P3a” component amplitude is responsive to stimulus probability, with larger responses to rare stimuli thought to reflect bottom-up prediction error signals that update context representations exogenously ^6,31^.

Considering the theoretical links between P300 subcomponents and perceptual decision-making efficiency, we hypothesised abnormal P300 responses (marking sub-optimal context updating) in patients with CRPS. Deficits at this level of decision-making could be potentially marked by either increases or decreases in P300 amplitude, with increases reflecting inefficient (resource-intensive) context updating, or decreases reflecting failed initiation of these mechanisms. It is also possible that hierarchically lower-level sensorimotor deficits could modulate P300 responses: such deficits may increase task difficulty and cognitive load and be reflected by augmented endogenous P3b responses, or may reduce bottom-up signalling of novel changes in sensory processing as indexed by the P3a component. Here, we sought evidence for these different possibilities by measuring SEPs elicited on stimulation of randomised digit locations across both hands in CRPS patients and matched healthy controls.

## Materials and Methods

### Study design and rationale

This was an experimental case-control study conducted in the EEG lab in the Herchel Smith Building for Brain and Mind Sciences, University of Cambridge, UK between March 2013 and July 2013. Ethical approval for the research was obtained by the East of England - Cambridge South ethics committee (reference number 12/EE/0305). The study was designed to detect group differences (between CRPS patients and healthy controls, HC) in mid-to-late latency (>100ms) tactile processing. In Experiment 1, participants performed a digit discrimination task to induce cognitive load and SEPs were investigated as a potential explanation for decrements in task performance. At the same time, the three middle digits on each hand were stimulated rarely compared to the outer digits in order to assess the effects of spatial probability. Experiment 2 had no task demands and had equiprobable digit stimulation, providing data on group differences independent of cognitive load. SEPs were recorded using high density (92-channel) EEG, making comparisons between groups, between CRPS-affected and unaffected sides of the body, and between digit types (high vs. low probability).

### Participants

Potential Complex Regional Pain Syndrome (CRPS) participants were identified from the CRPS UK registry and were approached for taking part in the study. Sample size considerations are in Supplementary Methods. The total number of potentially eligible CRPS patients contacted (who lived locally) was 30; 25 of these were confirmed eligible, of which 16 patients were able to be recruited before the recruitment period of the study ended. All patients were diagnosed with unilateral upper or lower limb CRPS (ruling out CRPS on the unaffected side) according to modified Budapest Research Criteria ^14^. The inclusion criteria were kept as broad as possible, including upper and lower limb-affected patients on the left or the right side. Although the study tests involved digit stimulation and discrimination on the hand only, previous work ^21^ has found that the location of CRPS symptoms (namely, upper vs. lower limb) does not significantly affect performance in digit discrimination, which is consistent with our hypothesis of deficits in hierarchically high-level decision-making mechanisms that are not somatotopically organised. Also recruited were 13 age-and-sex frequency-matched healthy (pain-free) volunteers. Healthy volunteers were recruited by advertising the study using posters in Addenbrooke’s Hospital, Cambridge, UK. All participants signed an informed consent form prior to taking part.

Data from 3 patients were excluded from the study analysis: one did not complete the study, and in the other two patients, data quality was extremely poor due to extreme movement artefact that could not be corrected or removed. This results in 13 patients (11 females, mean age=46.8 years). Of these patients, five had left arm, three had right arm and five had left leg affected respectively. The mean disease duration was 5.3 years (range 1-14). Demographic and medical details of the participants are shown in Supplementary Results (Tables S1 and S2). All participants were right handed, did not have any current or previous diagnosis of peripheral neuropathy, stroke, transient ischemic attack, multiple sclerosis, malignancy or seizure. The participants were required to refrain from consuming alcohol or smoking tobacco for 24 hours and caffeine for 12 hours prior to the study.

### Study Procedures

During the study visit, CRPS patients (but not healthy controls) were further characterised by using five questionnaires assessing pain severity, physical function, depersonalisation and mood: Brief Pain Inventory ^7^ – pain numerical rating scales only, Upper Extremity Functional Index ^37^, Lower Extremity Functional Index ^3^, Neglect-like Symptom Questionnaire ^13^, Hospital Anxiety and Depression Scale ^36^. Patients were not tested for the presence of referred sensations.

Participants were fitted with the EGI electrolyte cap with 128 channels (although only 92 channels were analysed – see pre-processing procedures). More details of the EEG set-up are in Supplementary Methods. Soft tactile stimuli were delivered to the tips of the digits of both hands using custom-made hand-boxes (one for each hand) that were calibrated to deliver non-painful stimuli with the same force. The participants were advised to report immediately if the sensation was uncomfortable or painful. The fingertips were also checked after each session to check for any redness of the skin. With this device and the EEG, two experiments were conducted as follows.

## Experiment 1

The main aim of Experiment 1 was to record (i) behavioural accuracy and response time for identification of each digit stimulated, (ii) SEPs related to task-relevant and spatially probabilistic tactile processing. The experiment consisted of 80 trials per block and four blocks per hand. The participants were given a small break of few minutes between each block. Only one hand was tested in a block and the order of the hand stimulation was randomised across blocks. Each trial lasted a maximum of 3 seconds, or terminated sooner depending on the speed of participant responses. The 80 trials per block were split into 30 each for thumb (digit 1) and little finger (digit 5) and the remaining 20 split between the remaining three fingers (Figure 1a). This resulted in a significantly higher probability of digits 1 and 5 (37.5% of the time for each, or 75% in total) being stimulated compared to digits 2, 3 and 4 (8.3% each, or 25% in total). Over the 4 blocks, 80 trials were presented for the total of the middle three digits (D2-D4) and 120 for each of the little finger and thumb; this provided more than enough data for robust measurement of the P300 potential, thought to require a minimum of 36 clean trials ^11^.

**Figure 1:**
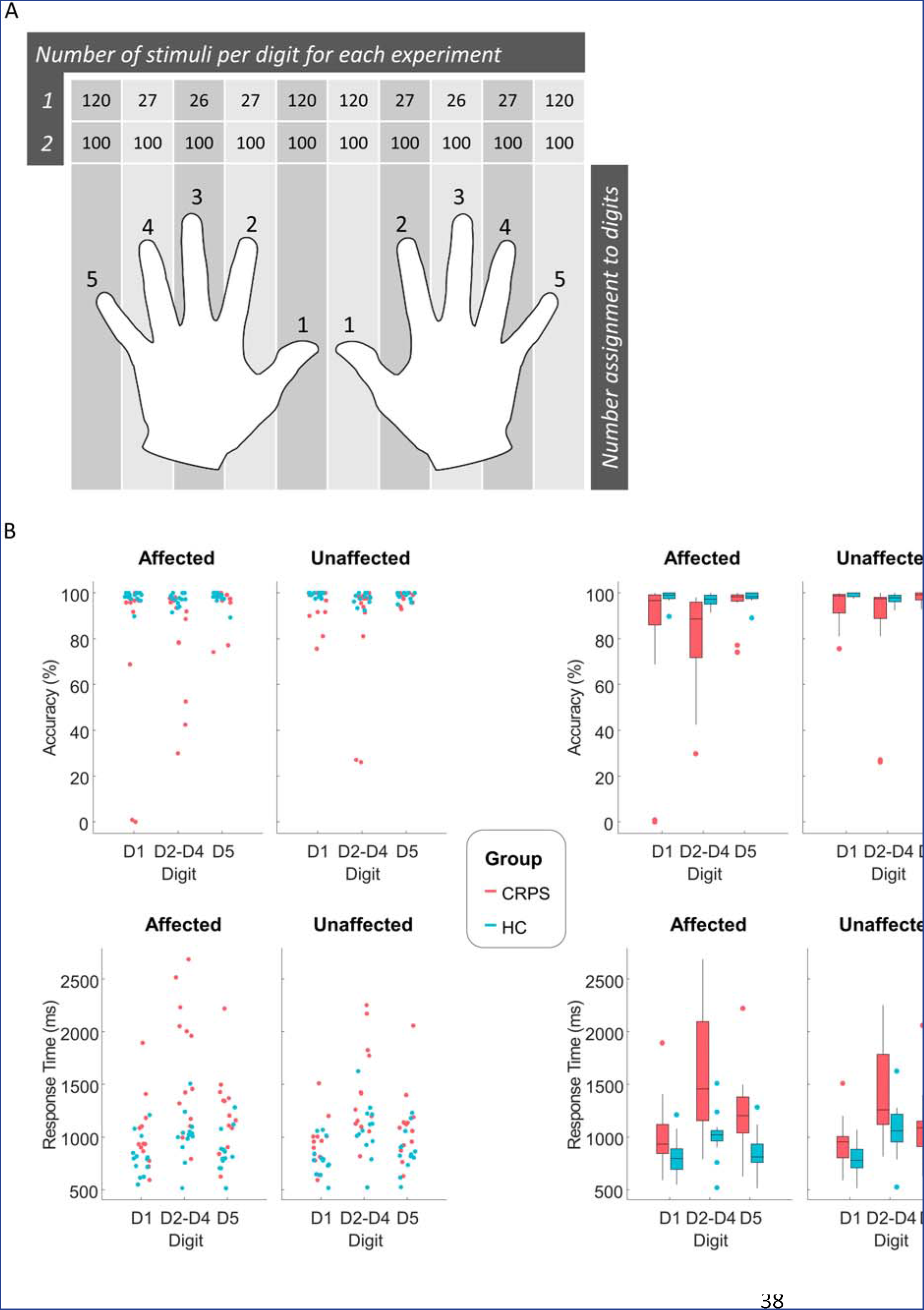
(A) The number of tactile stimuli delivered to each digit (randomised on each hand in turn) and the number assignment to each digit that the subject used to respond as to which digit was stimulated. (B) Behavioural results. Top: Accuracy data. Bottom: Response time data. Plots on the left (jittered dot plot) and right (box plot) are different representations of the same data.

For each block, participants were instructed to place one hand on the corresponding hand-box at a time. The digit positions on the hand-box were numbered consecutively from one to five starting with thumb (i.e.; thumb=1, index finger=2, middle finger =3, ring finger=4 and little finger=5) – see Figure 1a. Each time the subject received a stimulus (with duration of 0.05s) on a digit they responded by verbalising the number corresponding to that digit. A microphone attached to the EMG leads on the polygraph input box was used to capture the participant’s response time, measured as the time from the delivery of the stimulus to the start of the voice deflection on the EMG lead recording. Following verbal responses, the experimenter manually triggered the next stimulus by a key-press. The time taken for the experimenter to enter the response could not be strictly standardised but an effort was made for it to be as consistent as possible.

## Experiment 2

The main aim of the Experiment 2 was to study group differences in SEPs in the absence of cognitive task demands. In this experiment, participants were instructed to sit relaxed with eyes closed and head still and without responding to stimuli. 100 stimuli were delivered per digit of each hand (Figure 1a), split into two blocks per hand with the block order randomised. The digit order of stimulation was randomised within each block. Stimuli were delivered at 1 Hz: the stimulus duration was 0.05s with an inter-stimulus interval of 0.95s. There was a 10s break after every 50 stimuli.

### EEG data acquisition and pre-processing

During the experiment, 128-channel high-density EEG was recorded (but only 92 channels analysed) using the Net Amps 300 amplifier (Electrical Geodesics Inc., Oregon, USA). Due to the use of naturalistic touch stimuli with relatively long stimulus durations (compared to electrical stimuli, for example), early components (e.g. <100ms) were expected to be difficult to detect, and as our main interest was long-latency components, the sampling rate was set at 250 Hz. The vertex electrode (Cz) was used as a reference. Data from 92 channels over the scalp surface (at locations shown in supplementary Figure 1) were retained for further analysis, with channels excluded on the neck, cheeks and forehead, which mostly contributed movement-related noise than signal in patients.

EEG data pre-processing was performed using EEGLAB version 13.1.1 ^9^. Continuous data were initially high-pass filtered at 0.5 Hz and low pass filtered at 30Hz. After filtering, data were segmented into epochs including from 200ms preceding the stimulus to 800ms post-stimulus. Data containing excessive eye movement or muscular artefact were rejected by a quasi-automated procedure: noisy channels and epochs were identified by calculating their normalised variance and then manually rejected or retained by visual confirmation. Independent component analysis (ICA) based on the Infomax ICA algorithm ^2^ was run on the clean data excluding bad channels using the ‘runica’ function in EEGLAB. ICA components were visually inspected and bad components rejected. Bad channels previously identified by visual inspection were then replaced by spherical spline interpolation of neighbouring electrodes. Data were then re-referenced to the average of 92 channels. ERPs were calculated for each subject, experiment and digit type by averaging epochs.

### Definition of conditions for behavioural and ERP analyses

For all analyses, we considered three factors in the analysis: Digit Type, Side affected (by CRPS), and Group (CPRS, HC). Some explanation of these factors is provided here. The ‘Side’ factor levels were defined as affected and unaffected based on CRPS clinical assessment (Table S2). 4 patients were affected on the right side (2 upper limb, 2 lower limb) and 9 patients on the left side (5 upper limb, 4 lower limb). Due to the small sample size, we could not consider left-affected and right-affected CRPS patients, or upper and lower limb-affected patients, as separate groups. This has a disadvantage, in that two-thirds of patients were left-side affected, meaning that any comparison of affected and unaffected sides would be imbalanced with regard to whether left (non-dominant) or right (dominant) sides of the body were stimulated in each condition. To control for this, any inference of “Side” effects (i.e. affected vs. unaffected) in the CRPS patients were only interpreted with respect to any such effects in the HC group (e.g. by use of Group by Side interaction effects), and only after balancing the HC group data as follows: For the HC group to act as an adequate control, because no limbs were affected, we chose to consider four of the healthy participants’ right arm data and the remainder of the healthy participants’ left arm data as the ‘affected’ condition for the HC group, to match the left/right ratio of side affected in the CRPS group. The four HCs whose right-arm data was designated as ‘affected’ were chosen as those recruited in the same order as those CRPS patients who right arms were affected (i.e. participants 1, 4, 7 and 8 – Table S2). To be explicit, the term ‘affected’ in the HC group denotes a control condition for the CRPS ‘affected’ condition, and does not imply the presence of CRPS symptoms in the HC group.

In Experiment 1 only, for the ‘Digit Type’ factor, trials from stimulation of digits D2, D3 and D4 on each hand were considered as a single level which we subsequently label as ‘D2-D4’, and analysed alongside D1 and D5 to result in three digit types in total. Aside from their spatial location, D2-D4 digit trials were considerably more rare (low probability) than D1 or D5 trials (see Experiment 1 procedure). For Experiment 2, on the other hand, all digits were considered as separate levels (5 in total).

### Behavioural Data Analysis

Prior to testing our main hypothesis regarding cognitive cortical processing in CRPS, we sought to replicate and extend the results of previous work ^12,21^ that found reductions in accuracy and increases in response time in similar digit discrimination tasks in CRPS patients. Specifically, we tested a number of hypotheses, namely (H1) that there would be a group difference in at least one of these metrics in the same direction as found in previous studies, i.e. lower accuracy and longer response times, (H2) in the CRPS group (compared to the HC group), at last one of these metrics would indicate worse performance on the affected vs. the unaffected side of the body, indicating some somatotopic specificity, (H3) there would be slower response times to stimulation of rarer spatial locations (D2-D4) than more frequent locations (D1 and D5) due to the cognitive cost of switching attention to rarer locations, (H4) this cognitive switching cost would be less evident in the CRPS group compared to the HC group if they are less accurate at discriminating between digits in the first place. As these are all directional hypotheses, one-tailed statistics were used.

Behavioural data from Experiment 1 were analysed using IBM SPSS (Statistical Package for Social Sciences) software version 21 ^17^. Data for accuracy and response times were not normally distributed, especially in the CRPS group. Hence, non-parametric tests were performed to investigate overall group differences averaged over all conditions (Mann-Whitney U test), and to investigate within-subject condition effects in each group separately (Wilcoxon signed rank tests). Effect sizes are reported for non-parametric statistics using the formula *Z*/√*N*, where *Z* is the z-value output of the test and *N* is the number of samples in the test. This provides an effect size equivalent to the Pearson’s coefficient *r*, commonly interpreted as a small, medium and large effect with values of 0.1, 0.3 and 0.5 respectively.

### EEG Sensor-Level Analysis

SPM12 (www.fil.ion.ucl.ac.uk/spm) was used for subsequent EEG analysis steps ^22^. A “sensor-space” analysis was conducted (the standard approach implemented in SPM ^23^) and is described further in Supplementary Methods. For all analyses, the temporal window-of-interest was 100 – 400ms.

Analyses required direct comparisons between data derived from stimulation of CRPS-affected vs. CRPS-unaffected sides of the body. Due to the small sample size, we could not consider left-affected and right-affected CRPS patients as separate groups. Without further spatial transformation of the data, then, collapsing the data for “affected” and “unaffected” stimulation conditions across left and right-affected patients would have resulted in a mixture of left and right-hand stimulations respectively within each condition. As such, the identification of lateralised responses (e.g. the expected contralateral dominance for certain ERP components) would have proved problematic as both contralateral and ipsilateral data (from different patients) would have been mixed together within each condition. To resolve this problem, all sensor-space images representing cortical responses to right-hand stimulation were flipped on the x-axis. This resulted in images that represented (actual or apparent) left-sided stimulation, such that all contralateral differences resulting from hand stimulation on affected vs. unaffected sides would appear in the right hemisphere of the sensor-space images rather than being dispersed between both hemispheres.

Statistical analysis was performed on the resulting images with correction for multiple comparisons using random field theory to take into account the smoothness of the data on the scalp (Kilner & Friston, 2010). To do this, a General Linear Model (GLM) was estimated at the group level consisting of the between-subject factors Subject and Group (CRPS, HC), and the within-subject factors Side (affected, unaffected) and Digit (D1, D2-D4, and D5). F-contrasts were then constructed. Three effects were relevant to our hypothesis: (1) the main effect of Group, to test whether there are overall differences in cognitive processing in patients with CRPS, (2) the interaction of Group*Side, to test whether cognitive processing differences in CRPS are specific to the affected side, (3) the interaction of Group*Digit Type (in Experiment 1 only), to test whether any group effects depend upon the spatial probability of stimulation. For Experiment 2, the GLM was estimated in the same way and the contrasts were the main effect of Group and the Group*Side interaction.

### EEG Source-Level Analysis

Analysis at the source level was based on time windows defined from group differences on the sensor level. Canonical sensor locations were coregistered with the canonical head model in SPM. Lead field computation used a boundary element model (Phillips, Mattout, & Friston, 2007). The Bayesian source reconstruction method in SPM was used (Mattout, Phillips, Penny, Rugg, & Friston, 2006) with Multiple Spare Priors to estimate sources across the temporal window of interest (100 to 400 ms). Subsequently, source activity was averaged in each time window corresponding to the latency of significant effects of interest from the sensor-space analyses. Using F contrasts, significant differences were identified in source space and reported significant at a cluster-level significance of p (FWE) < 0.05 when considering statistical maps thresholded at *p* < 0.001.

### Brain-Behavioural Correlations

Neural correlates of behavioural outcomes in the patient group (namely, the hypothesised lower accuracy and slower response times) were investigated in source space. Regions-of-interest were identified as those clusters in the parietal and frontal lobe showing significant Group effects in the SPM contrasts, and were extracted from the SPMs as eigenvariates. To reduce the number of comparisons being made, and because activity from bilateral sources was highly multicollinear at midline locations including precuneus and SMA, where bilateral group differences were evident, sources from both clusters were averaged together prior to correlation analysis. Correlations using Spearman’s rank coefficient were conducted in the patient group only to understand behavioural variance within this group (also, accuracy scores in the control group were not variant enough to justify such analyses within the control group).

### Secondary (exploratory) correlation analyses

As clinical data were collected using a number of questionnaires (clinical pain ratings, anxiety depression, limb functioning, neglect-like symptoms), we performed exploratory analyses to test for correlations between these clinical variables and digit discrimination task-related (behavioural and EEG) outcomes. As well as testing against EEG source activity, we also explored clinical variable relationships with sensor-space clusters. Finally, we explored relationships between sensor-space clusters in the EEG data across the different latencies, as well as source-space clusters across latencies.

## Results

See Supplementary Results for details of the participant characteristics.

### Digit discrimination performance: Worse and more variable in CRPS patients

We hypothesised worse digit discrimination performance in the CRPS compared to the HC group in at least one of the metrics of accuracy and response time (RT). This hypothesis was supported. Accuracy in digit discrimination was lower on average for CRPS patients vs. healthy controls (HCs), but as shown in Figure 1b this was clearly driven by a small number of CRPS patients. The remainder of the CRPS group’s accuracy overlapped with that of the HCs, leading to a distribution of data strongly skewed towards high accuracy. Inferential statistics on the accuracy data provided a medium-to-large effect size in comparing the two groups (Table 1) with a p value of just over 0.01 (which remains significant after Bonferroni correction for the two tests – on accuracy and RT – used to address the hypothesis). For the RT data (Table 2 and Figure 1b) the group effect was stronger. In the CRPS group, while the distribution of the data was skewed towards longer RTs (Figure 1b), this did not appear to be strongly driven by such a small number of patients as for accuracy data. In addition, RTs showed overall greater variability in the CRPS group compared to the HC group. On average, RTs were longer in CRPS compared to HC participants with inferential statistics (Table 2) revealing a large effect size with strong statistical significance.

**Table 1:**
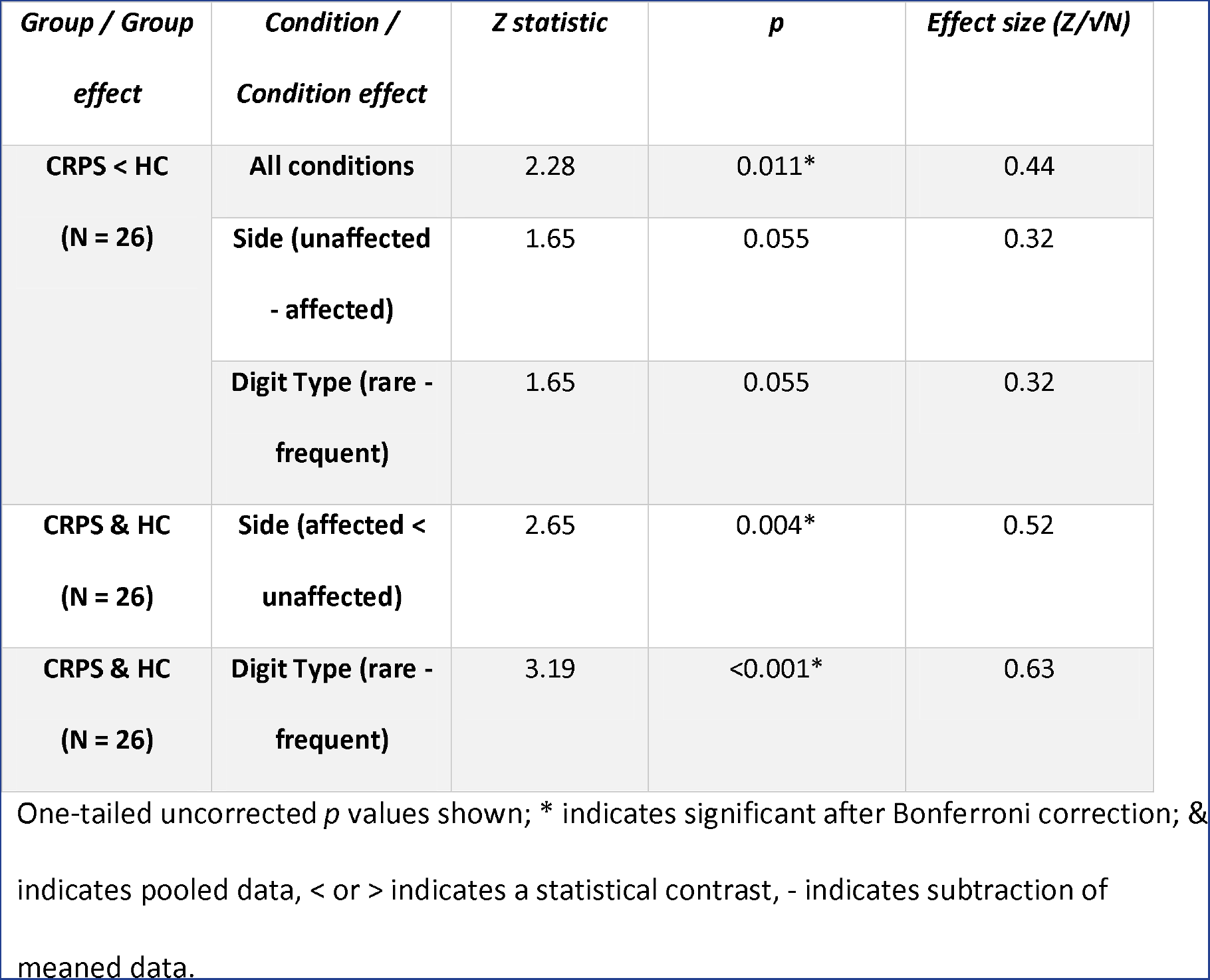
Non-parametric test statistics for accuracy

### Longer response times to CRPS affected vs. CRPS unaffected side stimulation

We further hypothesised that in the CRPS group, there would be worse performance (for at least one of the metrics) on the affected vs. the unaffected side of the body, indicating some somatotopic specificity in the CRPS group. As shown in Figure 1b, there was some indication of worse performance on the affected side compared to the unaffected side in CRPS patients. To address this hypothesis statistically, we looked at both the main effect of Side (pooling over both groups) and the interaction between Group and Side (a Group contrast on the subtracted Side data, i.e. unaffected minus affected side – Table 1). For accuracy data, there was a significant main effect of Side (remaining significant after Bonferroni correction for four tests) with a large effect size, but no significant interaction between Group and Side (although for this interaction there was a medium effect size with a p value of 0.055, suggesting caution is required in drawing strong conclusions from these results). Hence, there is little evidence for a Side effect on accuracy that is specific to the CRPS group, possibly due to a bias in performance in the HC group favouring the dominant hand (the “affected” condition in healthy controls mostly included data from the non-dominant hand). Conversely, for RTs there was no significant effect of Side across the two groups pooled together and the effect size was small (Table 2), but there was a significant interaction effect of Group and Side with a large effect size. It is visible from Figure 1b that on average, RTs were longer on the affected side in the CRPS group, but not in the HC group. Hence, RTs appear to be more specifically affected than accuracy by the side of the body stimulated in CRPS patients.

**Table 2:** Non-parametric test statistics for response time

| *Group / Group effect* | *Condition / Condition effect* | *Z statistic* | *p* | *Effect size (Z/√N)* |
| --- | --- | --- | --- | --- |
| **CRPS < HC**<br>**(N = 26)** | **All conditions** | 2.54 | 0.005* | 0.50 |
|  | **Side (unaffected - affected)** | 2.85 | 0.002* | 0.56 |
|  | **Digit Type (rare - frequent)** | 2.33 | 0.010* | 0.46 |
| **CRPS & HC**<br>**(N = 26)** | **Side (affected < unaffected)** | 1.41 | 0.080 | 0.28 |
| **CRPS & HC**<br>**(N = 26)** | **Digit Type (rare < frequent)** | 4.43 | <0.001* | 0.89 |

One-tailed uncorrected *p* values shown; \* indicates significant after Bonferroni correction; & indicates pooled data, < or > indicates a statistical contrast, - indicates subtraction of mean data.

### Performance decrements to rare stimuli are more apparent in CRPS patients

A separate hypothesis was that RTs would be longer on digits stimulated more rarely, but that this effect would be less apparent in the CRPS group due to a loss of discrimination accuracy. This hypothesis was only partly substantiated. A visible delay of RTs is evident for both groups in Figure 1b on more rarely stimulated middle digits (D2-D4) compared to more frequently stimulated digits (D1 and D5). Indeed, using inferential statistics pooling over both groups, there was a very strong Digit Type effect (Table 2). However, there was a medium-sized effect on the interaction between Group and Digit Type in the opposite direction to that hypothesised, with the mean difference in RTs between rare and frequently stimulated digits being 473 (SD 308) for the CRPS group and 203 (SD 108) for the HC group. In other words, there is no evidence to support the hypothesis that there is less cognitive cost to stimulating rare (vs. frequent) digit locations in CRPS patients because they are less able to discriminate the location changes.

There is also a suggestion from the boxplot of Figure 1b of lower accuracy on middle digits (D2-D4 condition), compared to D1 and D5, in the CRPS group. While this is not relevant to our hypothesis, for exploratory purposes, the statistics for the Digit Type effect and interaction between Group and Digit Type are also shown for the accuracy data in Table 1. In sum, accuracy was overall lower for rarer digit stimulations, but there was no evidence of this differing between groups.

### SEPs are augmented in CRPS patients at mid and late latencies

SPM sensor-space analyses were conducted on spatially-transformed sensor data (see Methods) such that all contralateral responses appeared on the right side of the scalp topography. In Experiment 1, the main effect of Group revealed significantly higher amplitude responses in the CRPS group compared to the HC group (Figures 2 and 3, and Table 3) in a contralateral negative-polarity cluster at 132ms (mean [95% CIs]: CRPS −2.65 [−3.26, −2.05]; HC −0.94 [−1.54, −0.34]) and a positive-polarity fronto-central cluster at 268ms (mean [95% CIs]: CRPS 4.34 [2.94, 5.73]; HC 2.08 [0.67, 3.47]). The positive peak at 268ms is consistent in timing and topography with a P300 response. However, there were no main effects of Digit Type or Side at either the 132ms or the 268ms latency. Exploring this further, this response was not qualitatively different when comparing the rarest (8% probability) to the less rare (37.5%) trials (Supplementary Figure S2). The lack of a Digit Type effect is contrary to our prediction that the P300 amplitude would be modulated by spatial probability.

**Figure 2:**
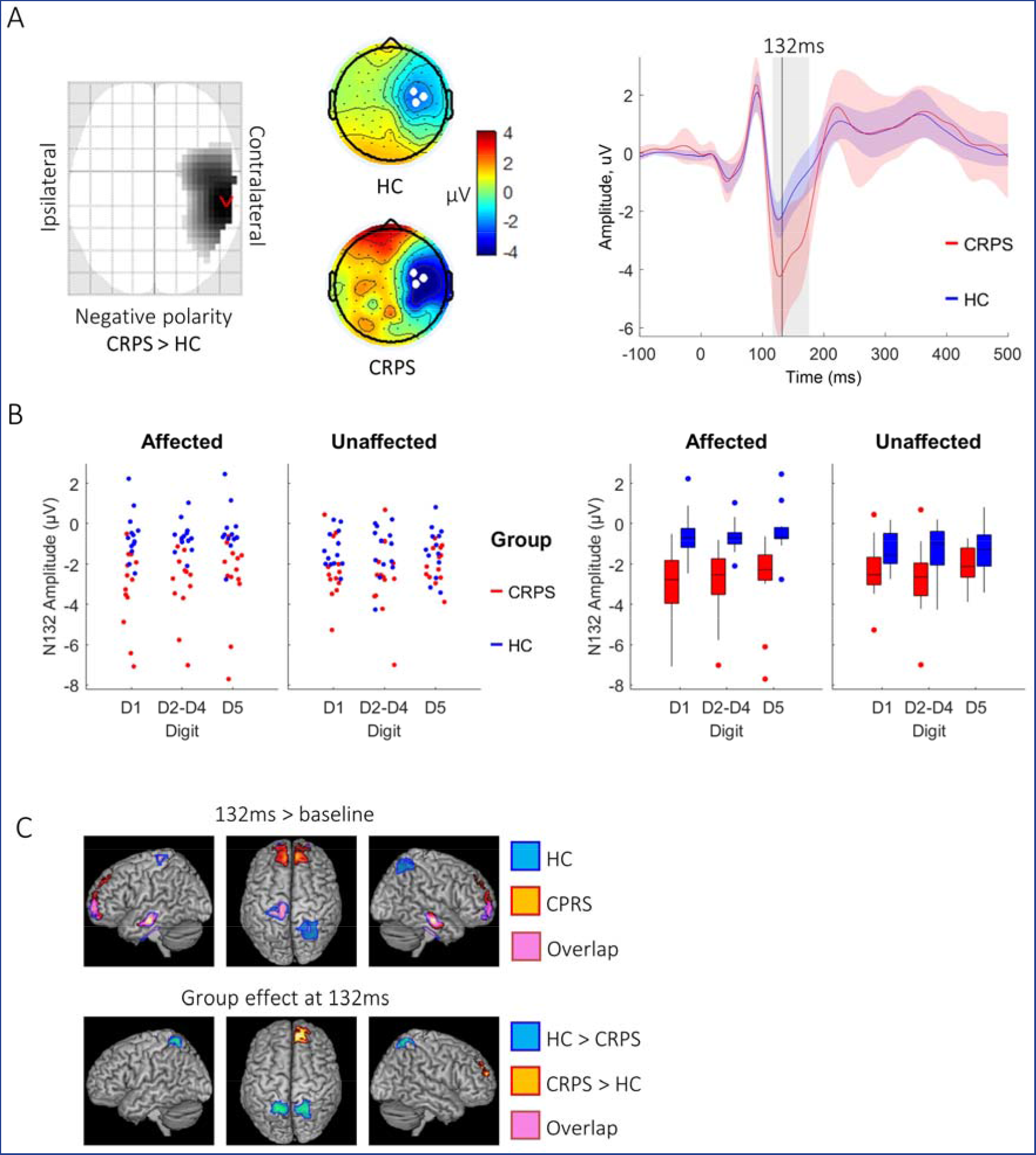
Experiment 1 data at 132ms. A) Left: Statistical Parametric Map identifying greater negative polarity response at 132ms in the CRPS vs. HC group over contralateral temporal and parietal electrodes. Image is thresholded at p < 0.001 uncorrected. Middle: Grand-average topographic maps from each group; white filled circles show the three electrodes with the most negative amplitude at 132ms that were averaged to display the waveform. Right: Waveforms with 95% confidence intervals showing the latency at 132ms (black line) and the temporal extent of the SPM sensor-space cluster (grey box). B) Two representations (jittered dot plot, left; box plot, right) of the activity in the significant sensor-space cluster for each group, side and digit type. C) Source-space results from the contrasts N132 activity vs. Baseline (-200ms to 0ms, top) and CRPS vs HC group (bottom) and their overlap (pink).

**Figure 3:**
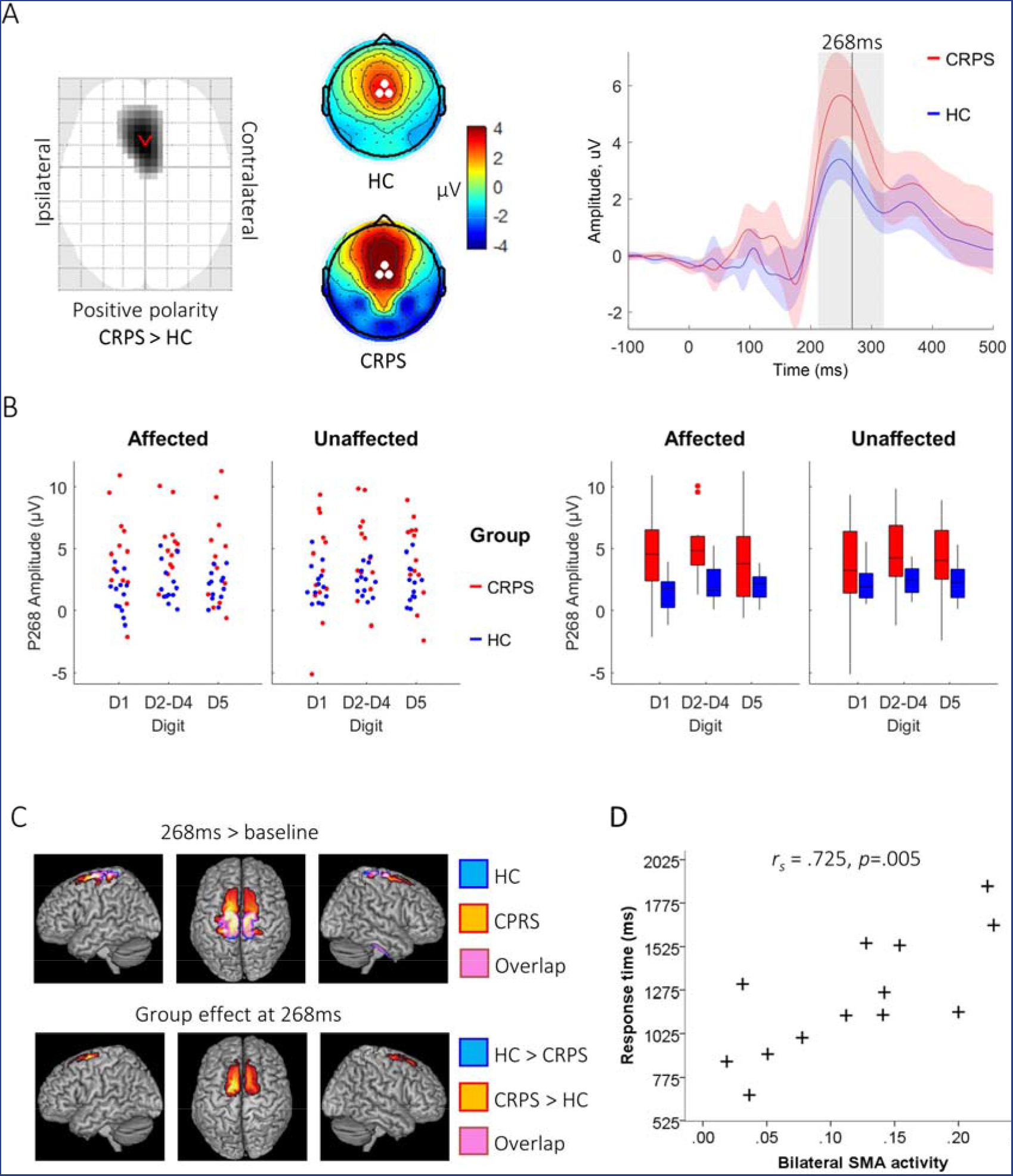
Experiment 1 data at 268ms. A) Left: Statistical Parametric Map identifying greater positive polarity response at 268ms in the CRPS vs. HC group over frontocentral electrodes. Image is thresholded at p < 0.001 uncorrected. Middle: Grand-average topographic maps from each group; white filled circles show the three electrodes with the most positive amplitude at 268ms that were averaged to display the waveform. Right: Waveforms with 95% confidence intervals showing the latency at 268ms (black line) and the temporal extent of the SPM sensor-space cluster (grey box). B) Two representations (jittered dot plot, left; box plot, right) of the activity in the significant sensor-space cluster for each group, side and digit type. C) Source-space results from the contrasts P268 activity vs. Baseline (-200ms to 0ms, top) and CRPS vs HC group (bottom). D) Non-parametric correlation of activity in bilateral Supplementary Motor Area (SMA, cluster defined from the CRPS > HC group contrast) with digit discrimination response time.

**Table 3:** Sensor-space results from the Group effect contrast

|  | **Spatial**<br>**coordinates**<br>**X, Y** | **Latency**<br>**(ms)** | **F** | **k** | **p FWE**<br>**(cluster-**<br>**level)** | **p FWE**<br>**(voxel-**<br>**level)** |
| --- | --- | --- | --- | --- | --- | --- |
| **Experiment 1** | 0, 18 | 268 | 34.31 | 2100 | 0.002 | <0.001 |
|  | 60, -30 | 132 | 26.76 | 1322 | 0.014 | 0.003 |
| **Experiment 2** | 21, -25 | 124 | 24.23 | 903 | 0.022 | 0.011 |

Spatial coordinates are indicated by X (left to right) and Y (posterior to anterior). Statistics include variance explained by the full model (*F* ratio), the cluster extent (*k*: number of voxels) and its associated cluster-level *P*-value (FWE corrected), in addition to the smallest voxel-level *P*-value (FWE-corrected) within the cluster.

At 132ms, the Group differences appear visually to be larger for the affected side compared to the unaffected side (Figure 2b); however, this did not translate into a statistically significant interaction between Group*Side, nor was there an interaction of Group*Digit Type. On the other hand, it is visually clear that at 268ms (Figure 3b) there are only Group effects and no interactions between Group and either Side or Digit Type, as borne out by the lack of statistical evidence for such interactions here either.

In Experiment 2, the main effect of Group revealed a similar contralateral effect as Experiment 1 although slightly earlier at 124ms (Figure 4a), with larger (more negative) responses in the CRPS group (mean [95% CIs]: CRPS −0.42 [−0.53, −0.31]; HC −0.18 [−0.29, − 0.06]). Again, mirroring the data from Experiment 1, there is the visual resemblance of a possibly larger Group effect for the affected side than for the unaffected side (Figure 4b), and yet no statistically significant interaction effects were evident. There was no Group effect at around the latency of a P300 response, but for comparison with Experiment 1 the data from Experiment 2 are plotted at 268ms (Figure 5).

**Figure 4:**
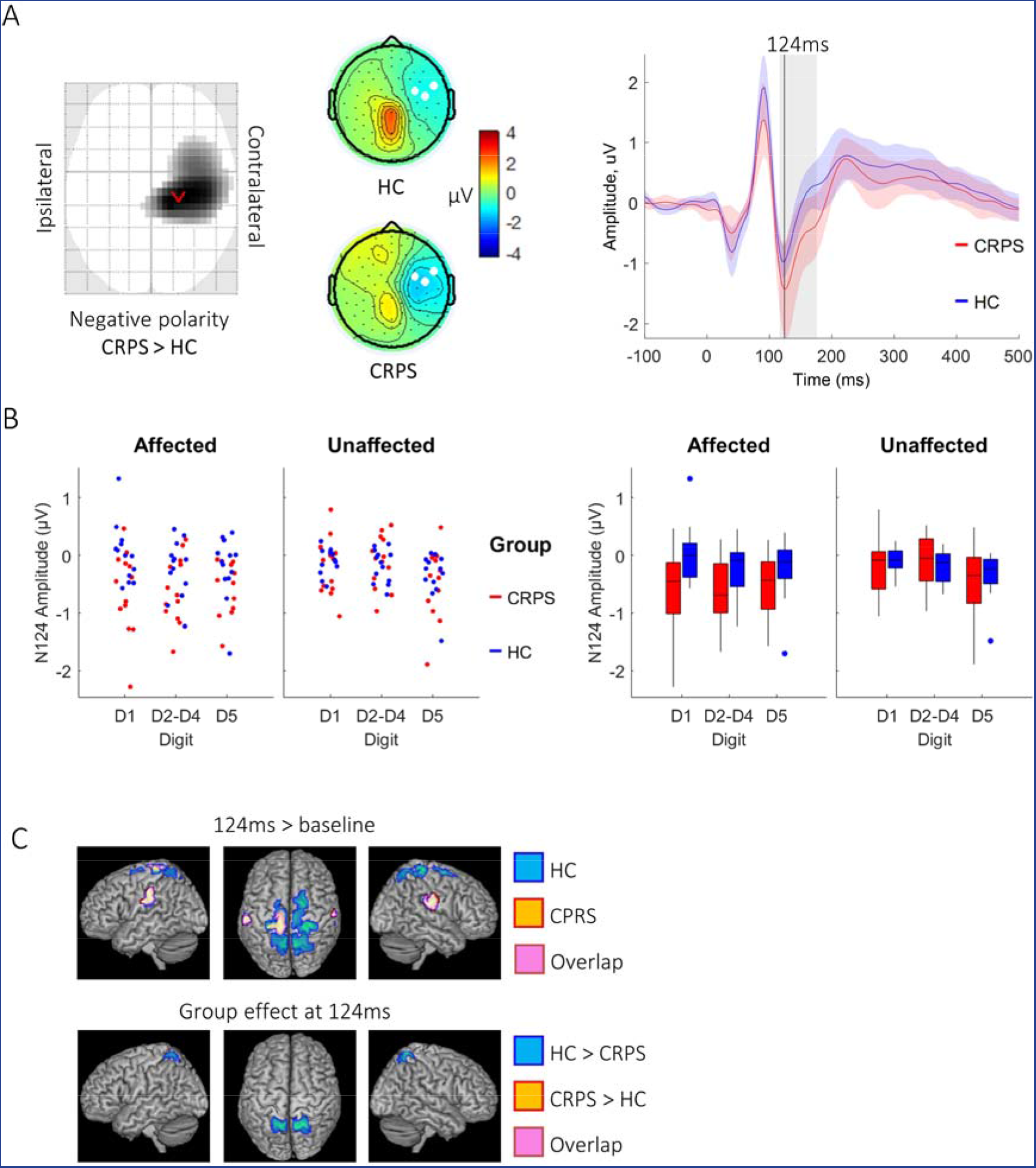
Experiment 2 data at 124ms. A) Left: Statistical Parametric Map identifying greater negative polarity response at 124ms in the CRPS vs. HC group over contralateral temporal and parietal electrodes. Image is thresholded at p < 0.001 uncorrected. Middle: Grand-average topographic maps from each group; white filled circles show the three electrodes with the most negative amplitude at 124ms that were averaged to display the waveform. Right: Waveforms with 95% confidence intervals showing the latency at 124ms (black line) and the temporal extent of the SPM sensor-space cluster (grey box). B) Two representations (jittered dot plot, left; box plot, right) of the activity in the significant sensor-space cluster for each group, side and digit type. C) Source-space results from the contrasts N124 activity vs. Baseline (-200ms to 0ms, top) and CRPS vs HC group (bottom).

**Figure 5:**
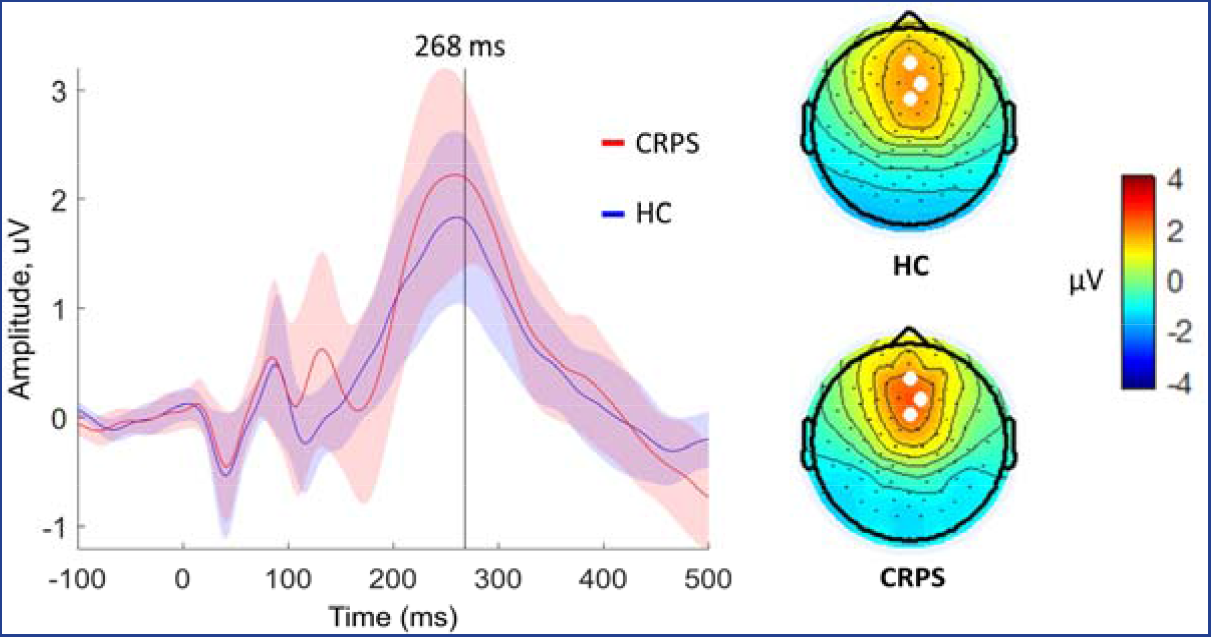
Experiment 2 data at 268ms demonstrating a P300-like response but no group effects. Left: Waveforms with 95% confidence intervals showing the latency at 268ms (black line) at which the topographic maps are displayed. Right: Grand-average topographic maps from each group; white filled circles show the three electrodes with the most positive amplitude at 268ms that were averaged to create the waveform.

### Reduced parietal and greater frontal cortical responses in CRPS patients

Sources of SEPs were estimated at latencies showing statistical significance in sensor-space. At each latency and for each group, contrasts are firstly reported for source activity (across all conditions) relative to pre-stimulus baseline (-200ms to 0ms) as one-sided t-contrasts (Tables S4 and S5, and Figures 2c, 3c and 4c). Secondly, group effects are reported as two-sided F-contrasts (Table 4). To summarise the results in relation to the hypothesised group differences, CRPS patients showed reduced activity in the precuneus and superior parietal lobe bilaterally in both Experiment 1 and Experiment 2 at middle latencies (132/124ms), suggesting a task-independent tactile processing deficit. On the other hand, greater activity was found in superior frontal areas in the CRPS group compared to the HC group in Experimental 1 (under task demands), which provides an explanation for the augmented responses found in the CRPS group in the analyses of sensor-space data. Specifically, middle latency responses were greater in a rostro-dorsal region of superior frontal lobe, while greater activity centred on the SMA at the later (268ms) latency. These results are discussed in greater detail in the Supplementary Results section.

**Table 4:** Source-space results for the contrast of CRPS group vs. HC group

|  | Latency<br>(ms) | Region | Spatial MNI<br>coordinates<br>X, Y, Z | k | P FWE<br>(cluster-level) | F | P FWE<br>(peak-<br>level) |
| --- | --- | --- | --- | --- | --- | --- | --- |
| Experiment 1 | 132 | Precuneus /<br>Sup. Parietal (I) | -22, -54, 66 | 268 | 0.004 | 24.15 | 0.04 |
|  | 132 | Precuneus /<br>Sup. Parietal (C) | 22, -56, 64 | 293 | 0.003 | 21.19 | 0.14 |
|  | 132 | Medial Sup. Frontal<br>(C) | 8, 52, 34 | 189 | 0.02 | 22.26 | 0.09 |
|  | 268 | SMA / Sup. Frontal<br>(I) | -16, 0, 68 | 764 | <0.001 | 57.83 | <0.001 |
|  | 268 | SMA / Sup. Frontal<br>(C) | 10, 4, 68 | 592 | <0.001 | 35.20 | <0.001 |
| Experiment 2 | 124 | Precuneus /<br>Sup. Parietal (I) | 14, -58, 66 | 469 | <0.001 | 29.08 | 0.012 |
|  | 124 | Precuneus /<br>Sup. Parietal (C) | -8, -56, 64 | 338 | 0.006 | 26.22 | 0.003 |

Spatial coordinates are indicated by X (left to right), Y (posterior to anterior) and Z (inferior to superior). Statistics include variance explained by the full model (*F* ratio), the cluster extent (*k*: number of voxels) and its associated cluster-level *P*-value (FWE corrected), in addition to voxel-level statistics including the minimum *P*-value (FWE-corrected) within the cluster.

### SMA source activity correlates with response time in CRPS patients

Temporal regions-of-interest from sensor-space topography maps (leftmost on Figures 2a, 3a and 4a) at 132ms and 268ms (Experiment 1) and 124ms (Experiment 2) were extracted for correlation analyses with behavioural data (accuracy and response time) in the patient group. No statistically significant correlations were found.

Regions-of-interest in the source data (clusters showing group differences in parietal and frontal areas) were also extracted for correlation analyses with behavioural data (accuracy and response time) in the patient group. Bilateral clusters were averaged together prior to analysis, resulting in a total of eight correlations (Table 5). After Bonferroni correction for multiple comparisons, a significant effect was found for response time - a positive correlation with source activity in SMA at 268ms during Experiment 1 (Figure 3d).

**Table 5:** Source-behaviour correlation results

|  | Experiment 1 |  |  | Experiment 2 |
| --- | --- | --- | --- | --- |
|  | Precuneus,<br>132ms | Sup. frontal,<br>132ms | SMA, 268ms | Precuneus,<br>124ms |
| **Accuracy** | $r_s = .033$ | $r_s = .418$ | $r_s = -.192$ | $r_s = -.484$ |
| | $p = .915$ | $p = .156$ | $p = .529$ | $p = .94$ |
| **Response time** | $r_s = .099$ | $r_s = -.060$ | $r_s = .725^*$ | $r_s = .412$ |
| | $p = .748$ | $p = .845$ | $p = .005$ | $p = .162$ |

*P* values are reported uncorrected; for multiple comparisons a threshold of significance is
applied of $p < 0.0063$ . Significant values are denoted by an asterisk.

### Late-latency SEPs predict physical functioning of the affected limb

Supplementary tables S6 to S8 show the results of exploratory correlation analyses. Due to the large number of comparisons in the correlations of SEP data with clinical variables, only one relationship merits attention due to its strong correlation (*r_s_* = .829, *p* = 0.0005) that survives multiple comparisons correction, namely the positive correlation between sensor-space cluster activity at 268ms in Experiment 1 with the lower/upper extremity function index, indicating that large P300-like responses predict higher levels of physical functioning of the affected limb in CRPS patients. Also of note is a correlation between the parietal and frontal sources showing group effects at 132ms in Experiment 1, which were correlated in both groups. Otherwise, sources were not correlated across latencies.

## Discussion

The findings are consistent with our main hypothesis that cognitive processing during tactile perceptual decision-making, downstream of early-latency somatotopic mapping, is altered in patients with CRPS. Secondarily, consistent with previous reports ^12,21^, there was evidence of less accurate digit discrimination and longer response times in CRPS patients compared to controls, but a high degree of between-subject variability indicating heterogeneity in the mechanisms driving these performance indices. Interestingly, response times appear to provide a more robust behavioural marker of perceptual decision-making deficits in CRPS patients than measures of digit discrimination accuracy: Not only was the response time difference between groups a stronger effect, but it was also more evenly distributed across patients and more sensitive to stimulation on the affected vs. the unaffected side of the body. Furthermore, response times provided utility in characterising sources of augmented P300-like responses in SMA, positively correlating with longer response times in CRPS patients. Conversely, decrements in activation of precuneus and superior parietal lobe in patients with CRPS (compared to healthy controls) were not related to performance markers; further investigation is required as to the origin and functional consequences of these parietal lobe decrements.

A limitation of the present study was the small sample size, which can impact the robustness and generalisability of group comparisons. Furthermore, the variable nature of the behavioural and EEG outcomes suggests that future research should focus on exploring this variability in relation to clinical heterogeneity. One example is disease duration (which in the current study ranged from 1 year to 14 years); to date, there has been no systematic investigation of abnormal tactile perception of cortical processing in CRPS in relation to this factor. A more complex confound is that of medication use: certain medications (such as opioids) may reduce cognitive performance, but also more severely affected patients are more likely to be prescribed such medication; establishing cause and effect is impossible from cross-sectional data. Lastly, it is instructive to note that digit misperceptions occur in other types of chronic pain ^21^. However, P300 responses from visual or auditory oddball stimuli can be either increased or decreased depending on the population of chronic pain patients studied ^18,40^. Further research is required to identify common and unique mechanisms driving these effects in patients with different diagnoses.

A further limitation is that the experiment failed to identify differential EEG responses to stimulation at more rare (8% probability) vs. less rare (37.5% probability) digit locations. It could be that the relative difference in rarity between conditions was not sufficient – 37.5% is still quite rare. On the other hand, response times were sensitive to the effect of rarity, and, counter to our predictions, CRPS patients were more sensitive to this effect rather than less. A plausible explanation is that longer response times on the middle three digits were affected by the additional spatial ambiguity arising from the fact that each of the middle digits (D2-D4) has two adjacent digits rather than just one for the outer digits (D1 and D5). CRPS patients may be more sensitive to this spatial ambiguity than controls. The experiment was therefore not well optimised to measure the effect of novelty-related responses as a potential marker of ineffective spatial discrimination in CRPS.

In the following, we discuss the results with reference to our two other predictions, that of augmented cognitive processing in patients specific to task demands, and altered task-independent processing related to spatial representations of the body.

### Late-latency SEPs mark abnormal cognitive processing in CRPS

A key finding relating to our hypothesis was the larger amplitude response at 268ms in CRPS patients compared to controls. The latency and scalp distribution of this effect is consistent with a P300; while also observed in Experiment 2, it was substantially diminished (indicating sensitivity to task demands) and there was no group difference. This supports the view that augmented task-related activity at 268ms in CRPS patients is due to greater attentional resource allocation required to perform the task.

However, the P300 is influenced by both endogenous (related to task demands) and exogenous (stimulus-driven) processes. The task-specificity of the group effect on the P300 suggests a role for endogenous processes, such that the high between-subject variance in the P300 response may relate to individual differences in perceived task difficulty. However, some characteristics of the response also bring us to cautiously suggest a role for bottom-up deficits in CRPS patients. There are three characteristics of our results that suggest the group difference at 268ms in the present study reflects differences in P3a-like novelty (exogenous) responses rather than P3b-like endogenous responses. Firstly, the early latency (for a P300) and fronto-central scalp distribution is reminiscent of a P3a ^8,31^. Secondly, the stimulus probabilities ranged from 8% to 37.5% depending on the digit, each of which can be considered spatially rare and therefore ‘novel’. Thirdly, previous studies of visual P3b-like components have identified that amplitudes are negatively related with RTs to the stimulus, possibly reflecting greater endogenous attention to the task ^34^; this is opposite to our finding in CRPS patients when considering SMA sources of the 268ms, a finding more consistent with stimulus-driven processes interrupting and delaying decision-making and subsequent response to stimuli. Shorter RTs to visual targets have also previously been found to be preceded by greater *pre*-stimulus activation of SMA, thought to promote top-down vigilance towards expected stimuli ^16^. In the present study, *post*-stimulus SMA activity predicted longer rather than shorter RTs, suggestive of stimulus-driven disruption of task performance rather than top-down vigilance. Further investigation is needed to test this hypothesis of specifically exogenous deficits in CRPS.

Interestingly, the magnitude of the P300 activity in CRPS patients predicted better limb functioning, suggesting it compensates to the disease rather than directly marking disease pathology, but we can only speculate about why this might be the case. While SMA is a premotor area that has an important role in the control of movements, it has also been implicated in broader perceptual and decision-making functions. Bilateral SMA (as well as, to a lesser extent, right precuneus) is particularly involved in maintenance of working memory required to discriminate between tactile stimuli separated by short delays ^15,32,35^, as well as in a subsequent decision-making phase ^33^. Therefore, it may be that higher-level cognitive processing of lower-level deficits in spatial representation of the body (e.g. in parietal lobe – see next section) in CRPS patients is adaptive in that it re-directs attentional resources and slows decision time in order to gather more sensory evidence to make more informed decisions. This explanation remains a cautious hypothesis until evidence is found that P300 amplitude variability in CRPS patients reflects an evidence accumulation process.

### Parietal lobe decrements in tactile processing in CRPS

Our analysis of SEPs and their sources at 124-132ms correspond to that commonly labelled as the N140 component of the SEP, which have been attributed to activity in SI and SII ^39,42^, as well as medial temporal (e.g. parahippocampal) regions, contralateral frontal cortex and insular cortex ^39,42^. Our source results are in broad agreement with this previous literature, except for additional activation of superior parietal lobe (SPL) and precuneus in our study.

In CRPS patients compared to healthy controls, we found a similar augmentation (in terms of latency and scalp distribution) of the negative polarity potential at 132ms in Experiment 1 compared to that at 124ms in Experiment 2. On the other hand, in both experiments, CRPS patients (compared to healthy controls) displayed decrements in tactile processing in the precuneus and SPL, which are likely related to the smaller positivity in posterior midline electrodes observable at mid-latencies. Hence, a mid-latency shift was observed in patients away from parietal processing and towards frontal activity. The fact that parietal lobe decrements were observed in both experiments shows that it occurs independently of task demands.

Functional imaging findings suggest the SPL and precuneus have an important role in visuo-spatial representation ^4^. Previously, research has found that precuneus is active during tracking of spatial changes in visual stimuli ^25^ and processes visuo-tactile mismatch responses in concert with the medial temporal lobe ^19^. Therefore, a plausible hypothesis is that reduced SPL and precuneus activity in tactile spatial tasks in patients with CRPS reflects deficits in visuo-spatial representations of the body. Behavioural characterisation of these cortical changes in future studies may be well served by measures of hand laterality recognition ^26^, on which performance is likely subserved by visuospatial functions.

### Conclusion

There is an unmet need to better characterise CRPS patients in terms of underlying mechanisms to aid early detection and treatment. Our study confirms highly variable tactile discrimination performance across CRPS patients, and points to previously undiscovered cortical processes at mid-to-late latencies relating to some aspects of task performance. Further research is required to fully characterise the pathophysiology and compensatory mechanisms underlying tactile discrimination performance in CRPS. Future studies would benefit from larger sample sizes that can cluster patients into mechanistically homogeneous subgroups and test for differential prognosis and treatment responses in these subgroups.

## References

1. Barrett LF, Simmons WK: Interoceptive predictions in the brain. Nat Rev Neurosci [Internet] 16:419–29, 2015 [cited 2016 Aug 9]. Available from: http://www.ncbi.nlm.nih.gov/pubmed/26016744

2. Bell AJ, Sejnowski TJ: An information-maximization approach to blind separation and blind deconvolution. Neural Comput 7:1129–59, 1995.

3. Binkley JM, Stratford PW, Lott SA, Riddle DL: The Lower Extremity Functional Scale (LEFS): Scale Development, Measurement Properties, and Clinical Application. Phys Ther 79:371–83, 1999.

4. Cavanna AE, Trimble MR: The precuneus: a review of its functional anatomy and behavioural correlates. Brain [Internet] Oxford University Press; 129:564–83, 2006 [cited 2017 Feb 9]. Available from: https://academic.oup.com/brain/article-lookup/doi/10.1093/brain/awl004

5. Chennu S, Finoia P, Kamau E, Monti MM, Allanson J, Pickard JD, Owen AM, Bekinschtein TA: Dissociable endogenous and exogenous attention in disorders of consciousness. NeuroImage Clin [Internet] 3:450–61, 2013 [cited 2017 Mar 24]. Available from: http://linkinghub.elsevier.com/retrieve/pii/S2213158213001381

6. Chennu S, Noreika V, Gueorguiev D, Blenkmann A, Kochen S, Ibáñez A, Owen AM, Bekinschtein TA: Expectation and attention in hierarchical auditory prediction. J Neurosci 33:11194–205, 2013.

7. Cleeland CS, Ryan KM: Pain assessment: global use of the Brief Pain Inventory. Ann Acad Med Singapore 23:129–38, 1994.

8. Comerchero MD, Polich J: P3a, perceptual distinctiveness, and stimulus modality. Brain Res Cogn Brain Res [Internet] 7:41–8, 1998 [cited 2017 Feb 14]. Available from: http://www.ncbi.nlm.nih.gov/pubmed/9714727

9. Delorme A, Makeig S: EEGLAB: an open source toolbox for analysis of single-trial EEG dynamics including independent component analysis. J Neurosci Methods 134:9–21, 2004.

10. DeMirci S, Savas S: The auditory event related potentials in episodic and chronic pain sufferers. Eur J pain 6:239–44, 2002.

11. Duncan CC, Barry RJ, Connolly JF, Fischer C, Michie PT, Näätänen R, Polich J, Reinvang I, Van Petten C: Event-related potentials in clinical research: Guidelines for eliciting, recording, and quantifying mismatch negativity, P300, and N400. Clin Neurophysiol [Internet] 120:1883–908, 2009 [cited 2017 May 15]. Available from: http://www.sciencedirect.com/science/article/pii/S1388245709005185

12. Förderreuther S, Sailer U, Straube A: Impaired self-perception of the hand in complex regional pain syndrome (CRPS). Pain [Internet] 110:756–61, 2004 [cited 2014 Sep 24]. Available from: http://www.ncbi.nlm.nih.gov/pubmed/15288417

13. Galer B, Jensen MP: Neglect-like symptoms in CRPS.Results of a self-administered survey. J Pain Symptom Manage 18:213–7, 1999.

14. Harden RN, Bruehl S, Stanton-Hicks M, Wilson PR: Proposed new diagnostic criteria for complex regional pain syndrome. Pain Med [Internet] 8:326–31, 2007 [cited 2014 Nov 11]. Available from: http://www.ncbi.nlm.nih.gov/pubmed/17610454

15. Hernández A, Zainos A, Romo R: Temporal evolution of a decision-making process in medial premotor cortex. Neuron 33:959–72, 2002.

16. Hinds O, Thompson TW, Ghosh S, Yoo JJ, Whitfield-Gabrieli S, Triantafyllou C, Gabrieli JDE: Roles of default-mode network and supplementary motor area in human vigilance performance: evidence from real-time fMRI. J Neurophysiol 109:, 2013.

17. IBM: IBM SPSS Statistics for Windows, Version 21.0. Armonk, NY: IBM Corp. 2012.

18. Karl A, Diers M, Flor H: P300-amplitudes in upper limb amputees with and without phantom limb pain in a visual oddball paradigm. Pain 110:40–8, 2004.

19. Kitada R, Sasaki AT, Okamoto Y, Kochiyama T, Sadato N: Role of the precuneus in the detection of incongruency between tactile and visual texture information: A functional MRI study. Neuropsychologia 64:252–62, 2014.

20. Kuttikat A, Noreika V, Shenker N, Chennu S, Bekinschtein T, Brown CA: Neurocognitive and Neuroplastic Mechanisms of Novel Clinical Signs in CRPS. Front Hum Neurosci [Internet] Frontiers; 10:16, 2016 [cited 2016 Aug 30]. Available from: http://journal.frontiersin.org/Article/10.3389/fnhum.2016.00016/abstract

21. Kuttikat A, Shaikh M, Oomatia A, Parker R, Shenker N: Novel Signs and their Clinical Utility in Diagnosing Complex Regional Pain Syndrome (CRPS) – A Prospective Observational Cohort Study. Clin J Pain [Internet] :1, 2016 [cited 2017 Feb 16]. Available from: http://www.ncbi.nlm.nih.gov/pubmed/27662180

22. Litvak V, Friston K: Electromagnetic source reconstruction for group studies. Neuroimage The Wellcome Trust Centre for Neuroimaging, University College London, UK.; 42:1490–8, 2008.

23. Litvak V, Mattout J, Kiebel S, Phillips C, Henson R, Kilner J, Barnes G, Oostenveld R, Daunizeau J, Flandin G, Penny W, Friston K: EEG and MEG data analysis in SPM8. Comput Intell Neurosci [Internet] Hindawi Publishing Corporation; 2011:852961, 2011 [cited 2017 Mar 22]. Available from: http://www.ncbi.nlm.nih.gov/pubmed/21437221

24. Marinus J, Moseley GL, Birklein F, Baron R, Maihöfner C, Kingery WS, van Hilten JJ: Clinical features and pathophysiology of CRPS. Lancet Neurol Elsevier Ltd; 10:637–48, 2011.

25. Meehan SK, Staines WR: The effect of task-relevance on primary somatosensory cortex during continuous sensory-guided movement in the presence of bimodal competition. Brain Res 1138:148–58, 2007.

26. Moseley GL: Why do people with complex regional pain syndrome take longer to recognize their affected hand? Neurology 62:2182–6, 2004.

27. Parra M a, Ascencio LL, Urquina HF, Manes F, Ibáñez AM: P300 and neuropsychological assessment in mild cognitive impairment and Alzheimer dementia. Front Neurol 3:172, 2012.

28. Di Pietro F, McAuley JH, Parkitny L, Lotze M, Wand BM, Moseley GL, Stanton TR: Primary somatosensory cortex function in complex regional pain syndrome: a systematic review and meta-analysis. J Pain [Internet] 14:1001–18, 2013 [cited 2014 Sep 11]. Available from: http://www.ncbi.nlm.nih.gov/pubmed/23726046

29. Di Pietro F, Stanton TR, Moseley GL, Lotze M, McAuley JH: Interhemispheric somatosensory differences in chronic pain reflect abnormality of the Healthy side. Hum Brain Mapp [Internet] 36:508–18, 2015 [cited 2015 Feb 4]. Available from: http://www.ncbi.nlm.nih.gov/pubmed/25256887

30. Pleger B, Ragert P, Schwenkreis P, Förster A-F, Wilimzig C, Dinse H, Nicolas V, Maier C, Tegenthoff M: Patterns of cortical reorganization parallel impaired tactile discrimination and pain intensity in complex regional pain syndrome. Neuroimage [Internet] 32:503–10, 2006 [cited 2014 Sep 11]. Available from: http://www.ncbi.nlm.nih.gov/pubmed/16753306

31. Polich J: Updating P300: An Integrative Theory of P3a and P3b. Clin Neurophysiol 118:2128–48, 2009.

32. Preuschhof C, Heekeren HR, Taskin B, Schubert T, Villringer A: Neural correlates of vibrotactile working memory in the human brain. J Neurosci 26:13231–9, 2006.

33. Preuschhof C, Heekeren HR, Taskin B, Schubert T, Villringer A: Neural correlates of vibrotactile working memory in the human brain. J Neurosci [Internet] 26:13231–9, 2006. Available from: http://eutils.ncbi.nlm.nih.gov/entrez/eutils/elink.fcgi?dbfrom=pubmed&id=17182773&retmode=ref&cmd=prlinks%5Cnpapers2://publication/doi/10.1523/JNEUROSCI.2767-06.2006

34. Ramchurn A, de Fockert JW, Mason L, Darling S, Bunce D: Intraindividual reaction time variability affects P300 amplitude rather than latency. Front Hum Neurosci [Internet] Frontiers Media SA; 8:557, 2014 [cited 2017 Feb 11]. Available from: http://www.ncbi.nlm.nih.gov/pubmed/25120458

35. Romo R, Salinas E: Flutter discrimination: neural codes, perception, memory and decision making. Nat Rev Neurosci [Internet] 4:203–18, 2003. Available from: http://www.ncbi.nlm.nih.gov/pubmed/12612633

36. Snaith RP: Health and Quality of Life Outcomes. 4:6–9, 2003.

37. Stratford PW, Binkley JM, Stratford D: Development and initial validation of the upper extremity functional index. Physiother Canada 53:259–67, 2001.

38. Tandon OP, Kumar A, Dhar D, Battacharya A: Event-related evoked potential responses (P 300) following epidural methylprednisolone therapy in chronic low back pain patients. Anaesthesia 52:1173–6, 1997.

39. Tarkka IM, Micheloyannis S, Stoki DS: Generators for human P300 elicited by somatosensory stimuli using multiple dipole source analysis. Neuroscience 75:275–87, 1996.

40. Tomasevic-Todorovic S, Boskovic K, Filipovic D, Milekic B, Grajic M, Hanna F: Auditory Event-Related P300 Potentials in Rheumatoid Arthritis Patients. Neurophysiology [Internet] Springer US; 47:138–43, 2015 [cited 2017 Sep 11]. Available from: http://link.springer.com/10.1007/s11062-015-9510-5

41. Turetsky BI, Calkins ME, Light GA, Olincy A, Radant AD, Swerdlow NR: Neurophysiological Endophenotypes of Schizophrenia: The Viability of Selected Candidate Measures. Schizophr Bull [Internet] 33:69–94, 2006 [cited 2017 Feb 13]. Available from: http://www.ncbi.nlm.nih.gov/pubmed/17135482

42. Valeriani M, Fraioli L, Ranghi F, Giaquinto S: Dipolar source modeling of the P300 event-related potential after somatosensory stimulation. Muscle and Nerve 24:1677–86, 2001.

43. van Velzen GAJ, Rombouts SARB, van Buchem MA, Marinus J, van Hilten JJ: Is the brain of complex regional pain syndrome patients truly different? Eur J Pain [Internet] 20:1622–33, 2016 [cited 2017 Sep 13]. Available from: http://doi.wiley.com/10.1002/ejp.882

